# Direction-Specific Effects of Biphasic Transcranial Magnetic Stimulation on Cortical and Cortico-spinal Excitability

**DOI:** 10.64898/2026.07.27.739487

**Authors:** Mirja Osnabruegge, Carolina Kanig, Wolfgang Mack, Berthold Langguth, Stefan Schoisswohl

## Abstract

**Aims & Methods:** Transcranial magnetic stimulation (TMS) is a well-established tool for inducing cortical excitation. However, the relevance of current direction on elicited effects is still incompletely understood. Combining TMS with electroencephalography (EEG) and electromyography (EMG) enables non-invasive analysis of evoked potentials both on cortical and peripheral level. In 23 healthy subjects, EEG and EMG responses to biphasic single pulses applied over the left motor cortex with anterior-posterior to posterior-anterior (AP-PA) or PA-AP current direction and 110% resting motor threshold (RMT) intensity were recorded and contrasted between the alternating phases. A cobot-assissted neuronavigation ensured stable coil-placement during the procedure.

**Results:** RMT was lower and EMG latency was shorter for AP-PA currents compared to PA-AP currents, whereas the EMG amplitude did not differ. For EEG responses, local and global evoked activity was higher for mid-components with PA-AP currents. P60 occurred earlier with PA-AP currents and N100 amplitude was higher in amplitude with AP-PA currents. The trial-wise MEP amplitude correlated significantly with P30 in the AP-PA and for both current directions with the N100 amplitude.

**Conclusion:** Our results highlight the directional sensitivity of M1 and the importance of further exploring the role of current direction in TMS protocols to better understand the cortical processes underlying cortico-cortical and cortico-spinal responses.

## Introduction

Transcranial magnetic stimulation (TMS) is a non-invasive brain stimulation method used for research, diagnostic and treatment purposes in health and disease. Via creation of a rapidly alternating magnetic field over the scalp, a current is induced in the cortex, which can depolarize superficial neurons. The magnetic field passes through the skull with little or no discomfort, which contributes to the good tolerability of the technique (Barker et al., 1985; Lefaucheur et al., 2020; Vucic et al., 2023). The combination of TMS with electroencephalography (EEG) and electromyography (EMG) allows to measure TMS-evoked cortical reactivity as well as to probe cortico-spinal activity.

By applying pulses of suprathreshold intensity over the contralateral motor cortex (M1), an evoked descending volley passes through cortico-spinal pathways on to the peripheral nerves. The evoked muscle activity respectively the motor evoked potential (MEP) is recorded by surface EMG attached to the target muscle (Barker et al., 1985; Di Lazzaro et al., 2004; Groppa et al., 2012). The required motor threshold, the amplitude of the evoked potential at defined suprathreshold intensities as well as the latency are all markers for the excitability of the motor system. Therefore, these output parameters can serve as indicators for deviant cortico-spinal tract excitability, which enables the use as potential markers for disease and treatment (Vucic et al., 2023).

With the simultaneous use of TMS and EEG, so-called transcranial evoked potentials (TEPs) can be recorded on the scalp surface, which reflect a direct readout for TMS-related cortical dynamics and provide information about cortical connectivity, excitability and brain state in a high temporal resolution to the stimulation (Bergmann et al., 2016; Ilmoniemi & Kicić, 2010; Tremblay et al., 2019). Applied over M1, the TMS pulse generates an immediate neuronal response under the coil, whose temporal and spatial course follows a typical pattern of deflections in the EEG with negative (N) and positive (P) peaks occurring at approximately 15 (N15), 30 (P30), 45 (N45), 60 (P60), 100 (N100) and 180 (P180) ms at different sensors (Ahn & Fröhlich, 2021; Bonato et al., 2006; Ilmoniemi et al., 1997; Schoisswohl et al., 2024; ter Braack et al., 2015). Source localization approaches depict the N15 as a direct activation in M1, whilst P30 is localized over ipsilateral premotor areas. N45, P60 and N100 occur over primary somatosensory areas and could reflect the afferent feedback of the peripheral motor system, while N100 and P180 could additionally be a marker for inhibitory neural activity or a result of sensory processing respectively sensory contamination (Ahn & Fröhlich, 2021; Schoisswohl et al., 2024; ter Braack et al., 2015).

However, the outcome parameters depend on technical parameters and in addition there is considerable individual variability in response to the same stimulation pattern (Pellegrini et al., 2018a; Silvanto & Pascual-Leone, 2008; Wassermann, 2002). Beside the influence of the stimulation intensity (Fox et al., 2006), parameters such as the width of the pulse, the waveform and the direction of the induced current flow affect the neuronal activation pattern (Casula et al., 2018; Di Lazzaro, Oliviero, Mazzone, et al., 2001; Di Lazzaro & Rothwell, 2014; Pellegrini et al., 2018b; Sommer et al., 2013). This may contribute to the heterogeneity and variability of findings on MEPs and TEPs and requires baseline comparative- and replication studies on the potential effects of the technical stimulation parameters to establish robust biomarkers.

Conventional TMS devices emit the voltage either in a asymmetric monophasic or symmetric cosine biphasic pulse shape (Neukirchinger et al., 2021). While for the monophasic shape only the initial phase is thought to induce stimulation, both phases of the biphasic pulse stimulate the underlying neuronal tissue, while the longer second phase is thought to be more effective (Groppa et al., 2012; Kammer et al., 2001).

Epidural recordings of M1 stimulation using monophasic pulses show that the direction of flow of the induced E-field plays a decisive role in MEP generation. An anteriorly directed posterior-anterior (PA) current flow with a coil handle at approximately 45°angle to the sagittal midline preferentially triggers indirect early volleys (I-waves) at lower intensities, which increase in number with ascending stimulation intensity (I_1_ - I_3_), resulting also in direct upstream activation (D-wave) at higher intensities (Di Lazzaro & Rothwell, 2014). Reversal of the current flow to anterior-posterior (AP) shows a more scattered activation pattern at corresponding intensities with a higher threshold, preferential activation of later I_3_-waves and a longer duration of the descending activation, resulting in longer MEP latencies but no differences in amplitude (Davila-Pérez et al., 2018; Delvendahl et al., 2014; Di Lazzaro, Oliviero, Saturno, et al., 2001; Di Lazzaro & Rothwell, 2014; Sakai et al., 1997; Sommer et al., 2006).

Biphasic pulses evoke a more inconsistent pattern of corticospinal volleys especially at low and threshold intensities. It partly follows monophasic patterns keeping in mind the more potent phase of the pulse is assumed to be the second one, e.g. applying the current in the tissue in an AP-PA direction results in lower active and resting thresholds plus shorter latencies and a trend-wise lower amplitude than with the PA-AP direction (Di Lazzaro et al., 2008; Guidali et al., 2023; Sommer et al., 2006; Weyh et al., 2005). By increasing the stimulation intensity, the recruitment pattern of the biphasic pulse becomes more similar to the pattern of the corresponding monophasic pulse (Di Lazzaro et al., 2008).

While the influence of current direction on MEPs has already been investigated in several studies, there are currently fewer findings for TMS-EEG recordings, particularly TEPs. However, these also suggest a directional sensitivity and preferential activation of different neuronal subsets akin to MEP findings.

At the level of TEP amplitude, subthreshold PA currents evoked higher positive amplitude peaks whereas negative amplitude peaks were more negative with AP and AP-PA currents (Casula et al., 2018). In addition, the early topographies differ between the current directions up to about 80 ms post-pulse in fronto-central regions. However, no difference in the local and global mean field power (LMFP, GMFP) between current directions were found but the RMT-adjusted stimulation intensities were higher for AP currents (Casula et al., 2018). The GMFP was first introduced by Lehmann & Skrandies (1980) and used by, for example, Esser et al. (2006) as a measure to characterize global EEG activity, while the LMFP refers to the activity only in region of interest, e.g., the motor cortex. Though, these results refer to activity evoked with shorter pulse widths than the pulse widths of currently available commercial devices, limiting comparability of evoked measures between studies employing different pulse settings (30, 80 µs vs. approx. 280 µs, Casula et al., 2018; Osnabruegge et al., 2024).

Focusing on the early P15 component, reflecting the contralateral positive peak over fronto-central areas and connectivity, Guidali and colleagues measured a smaller component amplitude and prolonged latency when using PA-AP currents in comparison to AP-PA currents (Guidali et al., 2023). Lucarelli and colleagues (2025) show on a descriptive level a higher, i.e. more negative N15, N45 and N100 amplitude with induced PA-AP currents. Concurrently, positive peaks of the P30, which was significant, P60 and P180 component were higher, i.e., more positive. Regarding the latencies of the TEP components, descriptively, the N15, P60, N100 and P180 occurred a few milliseconds later with PA-AP currents, while the P30 and N45 occurred earlier (Lucarelli et al., 2025).

In their study focusing on immediate responses (i-TEPs) Beck and colleagues demonstrated differences in the amplitudes of the three peaks occurring at approximately 2, 3 and 5 ms after TMS application depending on the current direction of the suprathreshold biphasic pulse. When adjusted to the respective RMT, the activation pattern of the early responses changed peak-wise, with AP-PA causing a larger amplitude in the first peak and smaller amplitudes in the second and third peak (Beck et al., 2024).

The described results also indicate a direction sensitivity of TEPs. However, further exploration of the influence of current direction on later TEP-components is necessary.

Within the framework of the present study, we want to address the necessity and examine the potential differences of transcranial evoked measures elicited with alternating current directions (AP-PA vs. PA-AP) of biphasic single TMS pulses applied to the motor cortex in healthy subjects. With the use of a collaborative robot (cobot) and neuronavigation, not only a precise coil positioning based on individual anatomical structures is guaranteed, but also potential coil-displacements due to e.g., head movement, that are automatically corrected while stimulation. Different responses in dependency of the current direction are expected: for MEPs, higher thresholds and longer latencies are anticipated in the PA-AP current direction but no definite direction for amplitude. For TEPs, respective higher positive and negative peak amplitudes as well as shorter latencies are expected to occur with AP-PA currents but no differences in the GMFP and LMFP between current directions are expected. Additionally, as both the TEP and the MEP are cortical excitability measures, we performed exploratory correlation analysis between cortical and peripheral potentials.

## Materials and Methods

The study was conducted at the University of Regensburg, Germany in accordance with the Declaration of Helsinki (ethic approval 21-2662-101). All participants gave written consent for participation and received monetary compensation.

### Participants

Participants were included if they were aged between 18-50 years, right-handed as assessed with the Edinburgh Handedness Questionnaire (EHI, Oldfield, 1971), had no contraindications to TMS and magnetic resonance imaging (MRI) (e.g. metallic implants, epilepsy or traumatic brain injury) as assessed by a physician, had no mental illness according to a screening questionnaire based on the DSM-V (American Psychiatric Association, 2013) and no depressive episode as assessed with the Major Depression Inventory (MDI, Bech et al., 2001). Participants were excluded if they did not fulfil one of the inclusion criteria, if they took medication or drugs altering brain function within the last six months prior to the experiment, if the resting motor threshold was higher than the security limits at the study centre of maximum 80% MSO or if they did not complete one measurement.

### Transcranial Magnetic Stimulation and Neuronavigation

A MagVenture X100 pulse source in combination with a butterfly figure-of-8 Cool-B65 A/P coil (both MagVenture, Farum, Denmark) was used to deliver biphasic pulses of AP-PA (PA-AP in coil) and PA-AP (AP-PA in coil) induced current directions with a jittered pulse interval of 10 ± 2 s and approximately 280 µs pulse width plus 500 ms recharge delay. For coil placement and stability, the TMS-collaborative coil holder (cobot, Axilum Robotics, Schiltigheim, France) in combination with the Localite neuronavigation system (Localite GmbH, Bonn, Germany) and the Polaris Spectra Camera (Polaris Spectra, NDI, Waterloo, Canada) were used. For individual coil positioning, a T1-weighted structural MRI (MPRAGE 160 slices, 256 × 256, voxel size = 0.977 × 0.977 × 1 mm3, flip angle 9◦, TR/TE/TI = 1910/3.67/1040 ms) was acquired from a 3 Tesla Siemens Scanner (Siemens, Munich, Germany) at the Department of Psychiatry and Psychotherapy, University of Regensburg, Germany. The cobot in combination with the individual MRI, neuronavigational software and tracking system allows a precise coil positioning and compensates for slow head movements of the participant, thus leads to a stable stimulation position during the measurement.

### Electrophysiological Data Acquisition

Electrophysiological data was recorded with the Brain Vision Recorder Version 1.25.0001 software with a BrainAmp DC system with BIP2AUX adapter in combination with solid gel Ag/AgCl Neuroline 710 surface electrodes (Ambu, Ballerup, Denmark) for EMG acquisition and 64-channel passive electrode caps for EEG acquisition (all Brain Products GmbH, Germany). Data was recorded at a sampling rate of 5 kHz.

### Measurement Procedure

The experiment consisted of one screening and one measurement session of approximately 2- and 3-hours duration. Within the screening session, participants were screened for inclusion and exclusion criteria, gave written consent and completed a structural MRI scan. To prepare the EEG and EMG, the scalp and hand area was prepared using Isopropyl alcohol 70% and abrasive paste (Nuprep, Weaver and Company, USA). The resistances were reduced to below 5 kΩ for scalp electrodes. EMG electrodes were attached to the FDI, ADM and APB in belly-tendon montage with the ground over the processus styloideus ulnaris. At the beginning of the measurement procedure, the hotspot and the resting motor threshold for both biphasic current directions (AP-PA and PA-AP) were determined using the semi-automated hotspot search method developed by our workgroup (Agboada et al. 2023). The neuronavigation system consisting of the Collaborative Coil Holder (Cobot, Axilum Robotics, Schiltigheim, France) and the Localite navigation software (Localite GmbH, Bonn, Germany) was utilized for precise control of stimulation target points in combination with subjects individual structural MRI images. Hotspot and RMT definition were followed by a stimulation block of 200 biphasic single pulses over the motor cortex at 110% RMT intensity. After a break of 10 minutes, a second stimulation block with the opposite current direction followed. The order of the current direction was randomized a priori. During measurement, participants wore ER-3C 10 Ω Insert Earphones (Etymotic Research Inc., USA) and ear muffs. Masking noise created via the TMS Adaptable Auditory Control (TAAC) toolbox (Russo et al., 2022) was delivered via an iPod Touch 7th Generation (Apple Inc., Cupertino, California, USA). The masking (70% white noise and 30% TMS click) loudness was adjusted to the individual sound perception level but not exceeding 85 dB SPL. The subjects were instructed to sit quietly, not to tense their hands, to look at a fixation cross on the opposite wall and to remove all electronic objects from their clothing and body.

### Preprocessing

Preprocessing of the TEPs was done with a custom written Matlab (Matlab 2022b; MathWorks, Natick, MA, USA) script, using the EEGLAB and Fieldtrip Toolboxes (Delorme & Makeig, 2004; Oostenveld et al., 2011; Rogasch et al., 2017), following the TESA (Rogasch et al., 2017) recommendations on EEG preprocessing. Additionally, to the TESA recommendations, a Fast-Fourier Transformation was used a priori to detect interfering harmonics and after identifying the pulse, an automatic removal of noise-contaminated channels was performed based on kurtosis with a z-score threshold of z = 4 and accompanied by visual inspection. Rejected channels were stored for later interpolation.

First, the channel locations were loaded and unused electrodes were removed. Triggers were identified in the C3 electrode. Data was then epoched in trials -300 to 500 ms around the TMS pulse and demeaned. The TMS artifact was removed -2 to 8 ms around the stimulus, and the missing data interpolated via cubic interpolation. After down-sampling from 5 kHz to 1 kHz, the trials were inspected visually and rejected if they contained corrupted data referring to eye blinks or movement during the time of stimulus application. Additionally, corrupted channels and the three channels with usual bad signal-to-noise ratio TP9, TP10 and Iz (Schoisswohl et al., 2024), were selected for rejection. The interpolated data at the time of the pulse was replaced with constant amplitude data. Next, the independent component analysis (ICA, *FastICA*) was applied to remove large amplitude eyeblinks or decay artifacts. Additionally, the source-estimate-utilizing noise-discarding (SOUND), and the signal space projection and source-informed reconstruction (SSP-SIR) algorithms were applied for artifact removal and noise reduction (Mutanen et al., 2018, 2022; Rogasch et al., 2014). The data was then band-pass filtered between 1 and 100 Hz and band-stop filtered between 48-52 Hz and 98-102 Hz with a 4^th^ order butter-worth filter. Data in the time range -2 to 8 ms around the TMS pulse was then again removed and interpolated via cubic interpolation. Missing channels were interpolated and the dataset re-referenced to average.

In the following step, the average 6 peak locations of each TEP component were used for the region of interest (ROI) selection per current direction and component within the pre-defined time of interest (TOI) of ± 5 ms for the early components N15, P30, N45 and P60 and ± 10 ms for the N100 and P180. Per subject, trial and component the peak amplitudes and latencies were extracted within the TOI and ROI. GMFP and LMFP potential were calculated using the following formula of Lehmann & Skrandies (1980) as used by Fieldtrip (Oostenveld et al., 2011) and e.g. Esser et al. (2006):

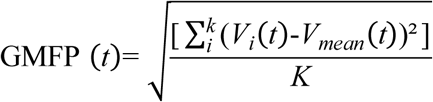

**Formula 1** – Global Mean Field Power calculation (Lehmann & Skrandies, 1980) where *K* equals the number of channels, *t* equals time and *V* equals the voltage at the channel *i*.

MEPs were preprocessed with a custom-written script and algorithm in Matlab (Matlab 2022b; MathWorks, Natick, MA, USA). Raw data was sampled at 5 kHz, epoched -300 to 500 ms around the TMS pulse and bandpass-filtered with 4^th^ order butter-worth 10 Hz high- and 500 Hz low-pass filter. If a priori interfering harmonics were detected with the Fast-Fourier-Transformation, a butter-worth 4^th^ order band-stop filter of ± 2 Hz was used. Peak-to-peak amplitude and latency were extracted in the time window 10-40 ms post-pulse. Latency was defined as that timepoint where 10% of the rising or falling flank of the first detected peak of the MEP after the TMS pulse in the time of interest occurred. Trials with artifacts (e.g., movement artifacts), a MEP amplitude ≤ 50 µV and an MEP-to-baseline ratio in the time window 200-50 ms pre pulse > 1.6 were automatically excluded by the custom-written algorithm.

### Statistical Analysis

All analysis were carried out via R (Version 4.3.3, The R Foundation for Statistical Computing with the packages *psych, rstatix, ggplot2, corrplot*) with the significance level set at 5% using non-parametric exact Wilcoxon signed-rank tests for contrasts between current directions. Non corrected *p*-values were interpreted and Bonferroni-corrected values (*p_adj_*) are reported for descriptive purposes only. Contrasts of interest were RMT, EMG amplitude, latency and EEG amplitude and latency. For explorative correlation analyses between peripheral and central measures of cortical excitability, only those trials that where neither rejected in the EMG or EEG preprocessing process were used for calculation of Spearman’s rank correlation coefficient.

## Results

### Sample

The final sample consisted of 23 participants with a mean age of 25 ± 5 years (14 females). Results of the questionnaires indicate a right-handed, non-depressed sample with a slightly above average vocabulary-based IQ (MDI score of 5 ± 3; MWT-B IQ 110 ± 12, Lehrl, 1989; EHI 86 ± 13). Two participants were excluded due to contraindications for the experimental procedure. For MEP analysis, two subjects had to be excluded due to insufficient signal-to-noise ratio in the EMG.

### EEG Analysis

#### Transcranial Evoked Potentials and Topographies

Exact Wilcoxon signed-rank tests for TEP components amplitude and latency showed a significant difference between current directions for the P60 latency, which occurred earlier with PA-AP currents (*p* = .03267, *p_adj_* = .4900) and for the N100 amplitude, which was higher with PA-AP currents (*p* = .009146, *p_adj_* = .1372). No other significant differences for TEP components were found (see **Figure 1 & 2** or Appendix for *p*-values of other comparisons). To visualize the data quality, **Figure 3** depicts the mean butterfly plots of all EEG channels as mean time course over all trials and subjects for both current directions. The topographical distribution shows an average peak activity at approximately 14 ms over left central and parietal areas for the N15 (10-20 ms) component and at approx. 30 ms over central and frontal regions for the P30 (25-35 ms) component. For the N45 (40-50 ms) component, the peak occurred at approx. 46 ms over central and right parietal regions, for the P60 (55-65 ms) at approx. 59 ms over left parietal and occipital areas. The later components N100 and P180 showed peak activations over central and left parietal sensors. In **Table 1**, the exact average amplitude and latency values and electrode clusters are listed per component.

**Figure 1.**
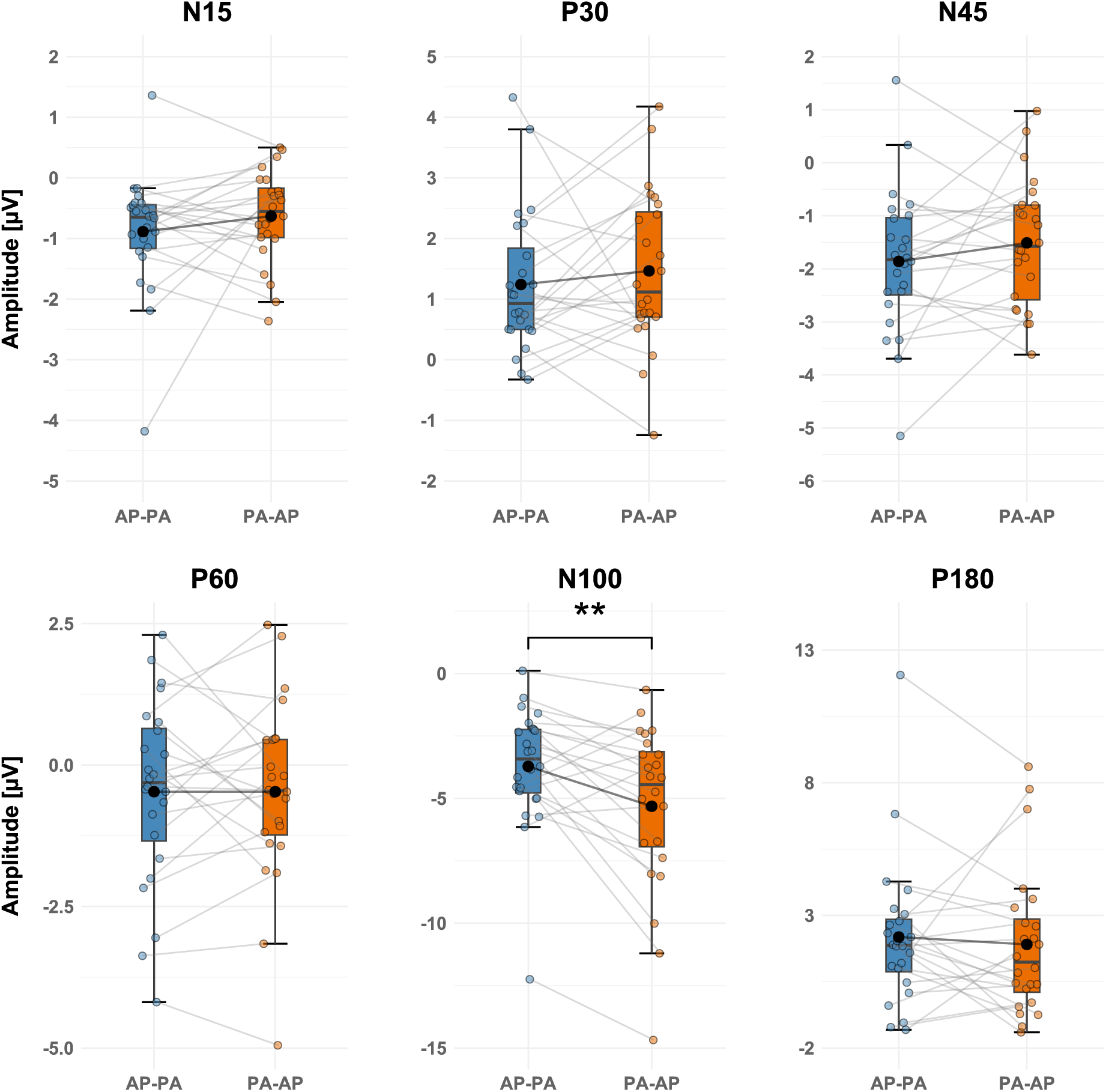
Comparison between Current Directions: Amplitude of Transcranial Evoked Potentials. Tukey Boxplots showing the experimental results for the TEP amplitude per component and current direction. Asterisks indicate significance levels of exact Wilcoxon signed-rank comparisons: \*\**p* ≤ .01. The single points represent the individual values per subject, connected by a grey line. The mean values per condition are shown in black. Whiskers depict the 25th percentile value minus 1.5 times the inter-quartile range and the 75th percentile plus 1.5 times the inter-quartile range. A detailed description of mean and standard deviation values can be found in Table 1.

**Figure 2.**
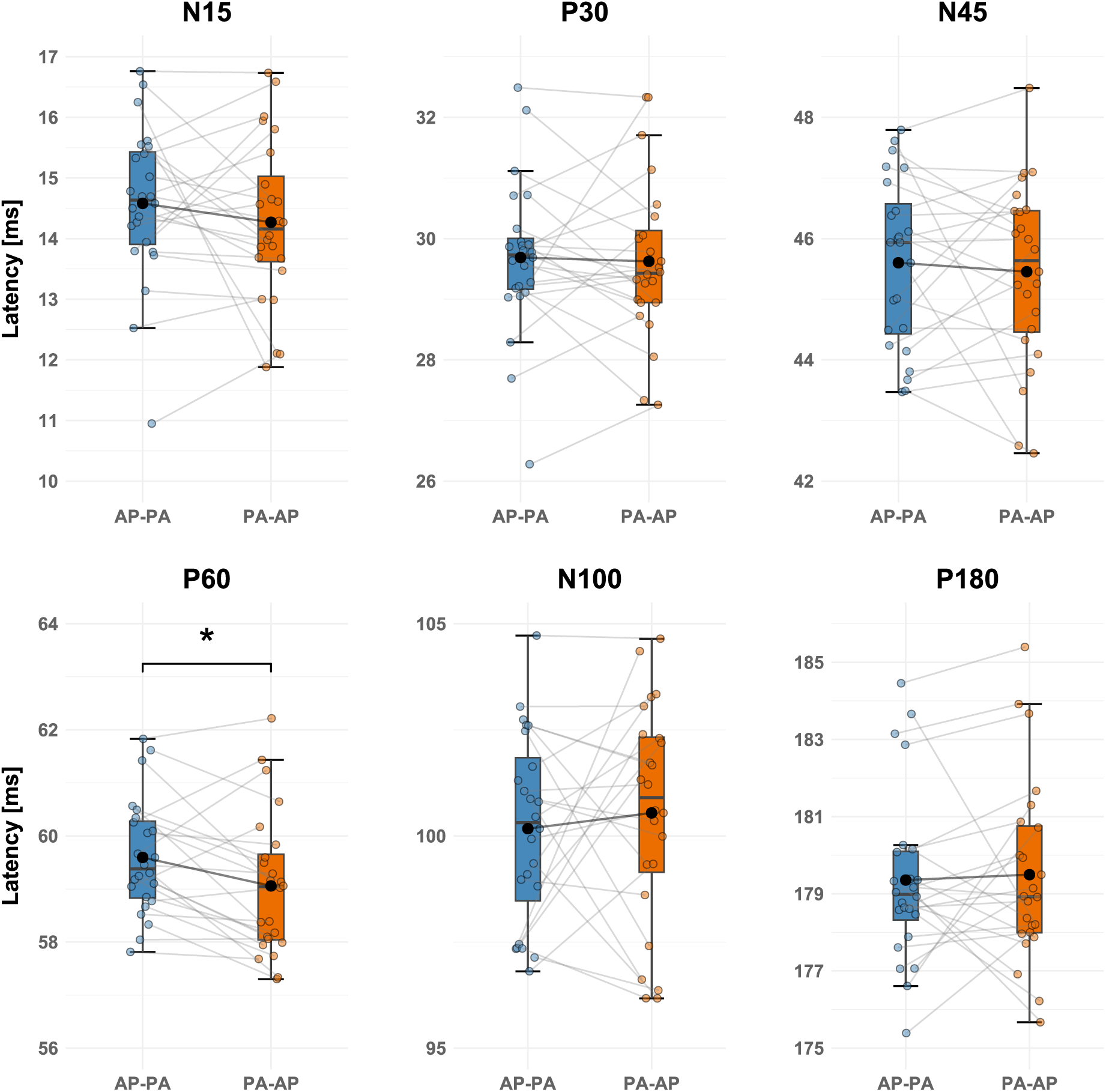
Comparison between Current Directions: Latency of Transcranial Evoked Potentials. Tukey Boxplots showing the experimental results for the TEP latency per component and current direction. Asterisks indicate significance levels of exact Wilcoxon signed-rank comparisons: \**p* ≤ .05. The single points represent the individual values per subject, connected by a grey line. The mean values per condition are shown in black. Whiskers depict the 25th percentile value minus 1.5 times the inter-quartile range and the 75th percentile plus 1.5 times the inter-quartile range. A detailed description of mean and standard deviation values can be found in Table 1.

**Figure 3.**
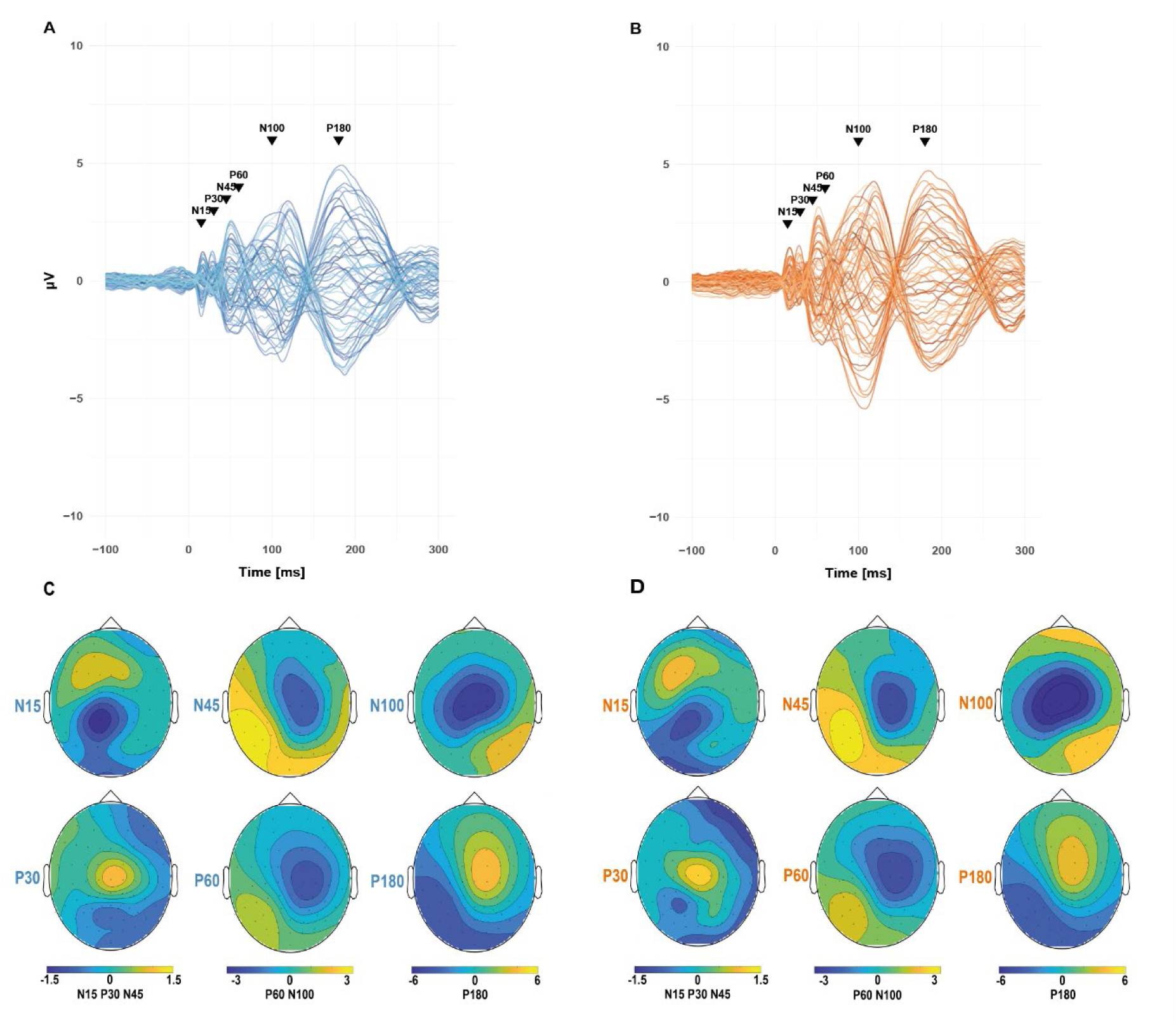
Butterfly Plots and Peak Topographies per Current Direction. A: Butterfly plot of the EEG trace for the AP-PA condition, averaged over all subjects, trials and electrodes. B: Butterfly plot of the EEG trace for the PA-AP condition, averaged over all subjects and trials and electrodes. C: Topographies of the peak activations per component and current direction. Upper row, from left to right: N15, N45 and N100. Bottom row, from left to right: P30, P60 and P180 elicited with AP-PA current direction. D: Topographies of the peak activations per component and current direction. Upper row, from left to right: N15, N45 and N100. Bottom row, from left to right: P30, P60 and P180 elicited with AP-PA current direction. Note the scales differ between components.

**Table 1.** Peak Component Amplitude, Latency and Electrode Clusters.

AP-PA: anterior-posterior to posterior-anterior current flow in the tissue. PA-AP: posterior-anterior to anterior-posterior current flow in the tissue. SD: standard deviation.
| | Amplitude [μV] | | Latency [ms] | | Peak Electrode Cluster | |
| --- | --- | --- | --- | --- | --- | --- |
|  | AP-PA | PA-AP | AP-PA | PA-AP | AP-PA | PA-AP |
| | Mean ± SD | | | | | |
| N15 | -0.89 ± 1 | -0.63 ± 0.78 | 14.58 ± 1.31 | 14.27 ± 1.38 | CP1 P1 CP3<br>Cpz P3 C1 | CP1 P3 P1 Iz<br>P5 CPz |
| P30 | 1.24 ± 1.2 | 1.46 ± 1.32 | 29.69 ± 1.31 | 29.62 ± 1.35 | Cz C2 C1 FC2<br>FC1 F7 | Cz C1 C2 FC1<br>FC2 FC3 |
| N45 | -1.86 ± 1.41 | -1.51 ± 1.23 | 45.6 ± 1.42 | 45.45 ± 1.53 | C2 FC2 CP2 C4<br>Cz CP4 | C2 C4 CP2<br>FC2 CP4 FC4 |
| P60 | -0.47 ± 1.68 | -0.47 ± 1.65 | 59.59 ± 1.11 | 59.06 ± 1.37 | P7 P5 PO7 TP7<br>CP5 O1 | P5 PO7 P7 P3<br>PO3 CP5 |
| N100 | -3.72 ± 2.5 | -5.32 ± 3.4 | 100.17 ± 2.29 | 100.54 ± 2.64 | Cz C1 FC2 C2<br>FC1 CP1 | Cz C1 FC2 C2<br>FC1 CP1 |
| P180 | 2.18 ± 2.87 | 1.91 ± 2.8 | 179.36 ± 2.29 | 179.49 ± 2.45 | FC2 C2 Cz F2<br>FC4 Fz | FC2 C2 F2 Cz<br>Fz FC4 |

#### Local & Global Mean Field Power

After splitting LMFP and GMFP into three blocks of early (10-50 ms), mid (51-110 ms) and late (111-200 ms) responses and contrasting them between current directions, mean local and global activity was significantly higher for mid components with PA-AP currents (LMFP: MD_PA-AP_ = 0.77, MD_AP-PA_ = 1.2, V = 73, CI [-0.567 -0.001], *p* = .04844, *p_adj_* = .1453; GMFP: MD_PA-AP_ = 1.7, MD_AP-PA_ = 2.38, V = 69, CI [-0.928 -0.022], *p* = .03543, *p_adj_* = .1063) but not for early and late components (*p* ≥ .1). The trial- and subject-wise averaged time course of the LMFP & GMFP is depicted in **Figure 4**.

**Figure 4.**
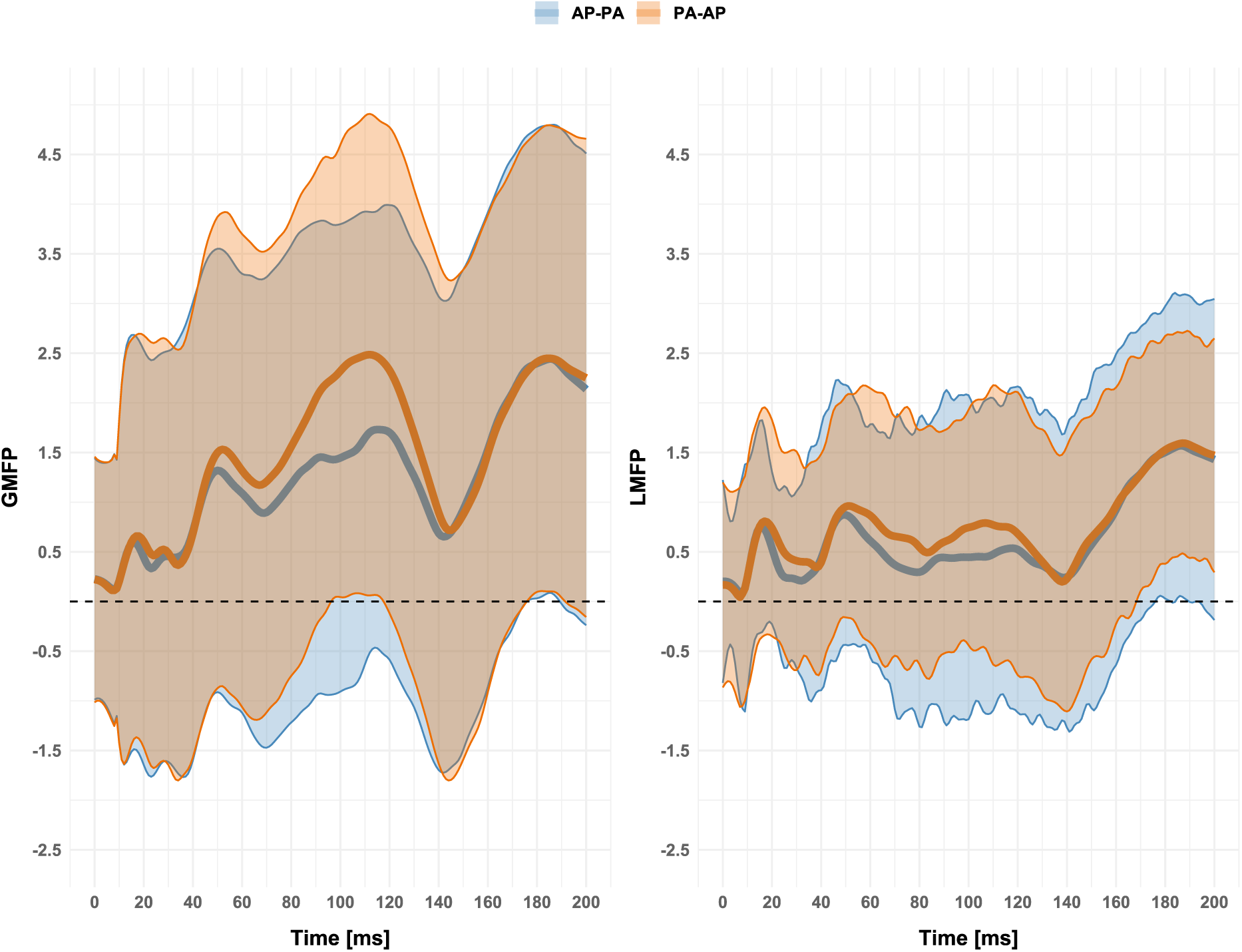
Comparison between Current Directions: Local- and Global Mean Field Power. Local- (LMFP) & Global Mean Field Power (GMFP) per current direction for the time range 0-200 ms post TMS-pulse ± SD, represented by the shaded area. LMFP was computed for the ROI-cluster with the electrodes FC1, FC3, C1, C3, CP1, CP3.

### EMG Analysis

Results of the EMG analysis are depicted in **Figure 5**. Exact Wilcoxon signed-rank test showed a significantly lower RMT with AP-PA currents (*p* ≤ .001, *p_adj_*= .0016). The latency of the MEP was significantly longer with PA-AP currents (*p* = .02901, *p_adj_* = .4352). There was no difference in MEP amplitude (*p* = .7079, *p_adj_* = 1). Detailed values and results of the explorative correlation analysis can be found in the Appendix.

**Figure 5.**
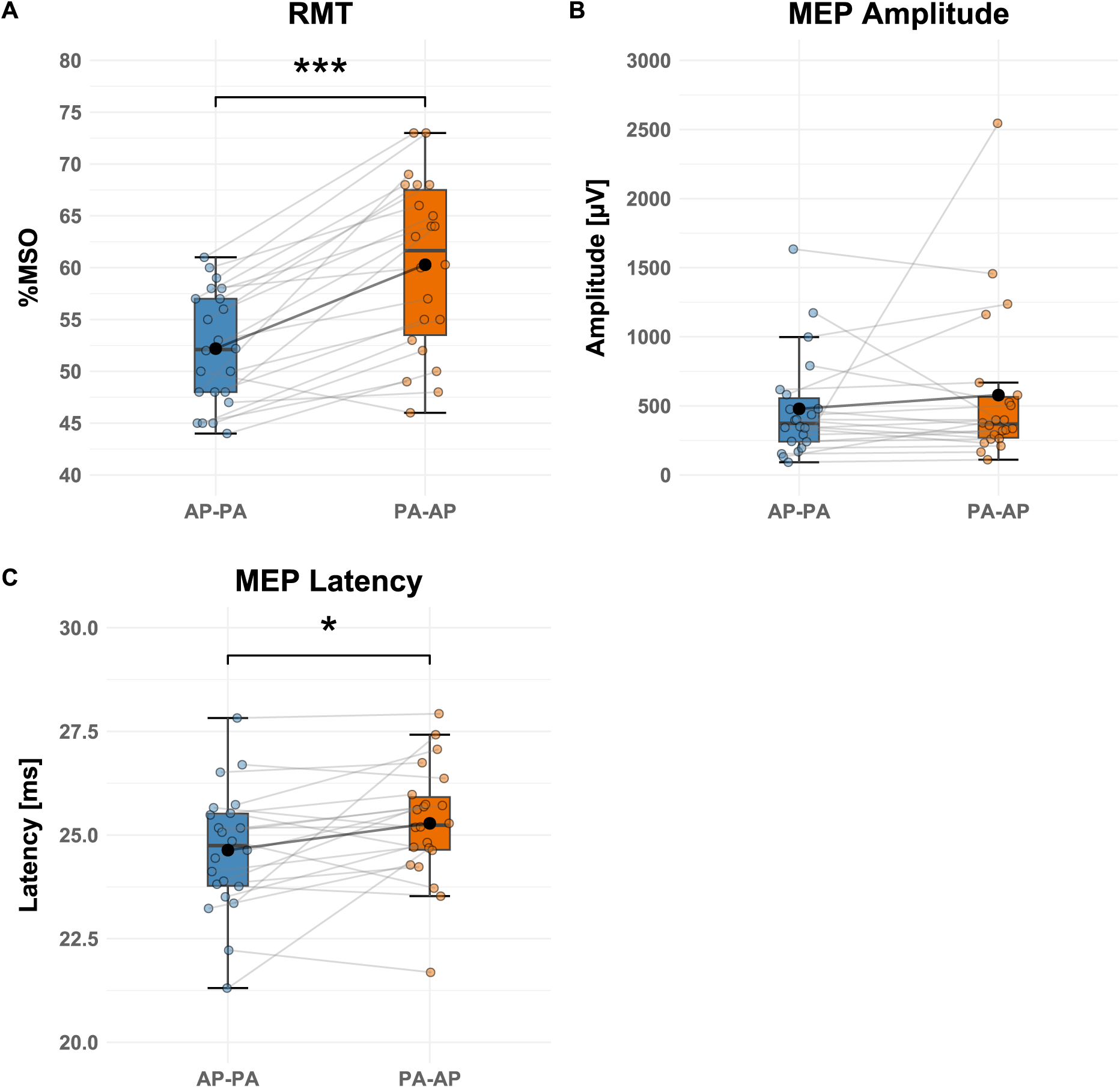
Comparison between Current Directions: Resting Motor Threshold and Motor Evoked Potentials. Tukey Boxplots showing the experimental results for the resting motor threshold (RMT, A), motor evoked potential (MEP) amplitude (B) and MEP latency (C) per current direction for 23 subjects. Asterisks indicate significance levels of the exact Wilcoxon signed-rank test \**p* ≤ .05, \*\*\**p* ≤ .001. The single points represent the individual values per subject, connected by a grey line. The mean values per condition are shown in black. Whiskers depict the 25^th^ percentile value minus 1.5 times the inter-quartile range and the 75^th^ percentile plus 1.5 times the inter-quartile range. A detailed description of median, mean and standard deviation values can be found in the Appendix.

### Correlation Analysis

Spearman rank correlations showed significant positive correlations between trial-wise P30 and MEP amplitude with AP-PA (ρ = 0.1589, *p* = .0237, *p_adj_* = .119) and between N100 and MEP amplitude for both current directions (ρ_AP-PA_ = -0.2268, *p* = .0012, *p_adj_* = .0062; ρ_PA-AP_ = -0.1876, *p* = .0078, *p_adj_* = .0390) on the group-level. Significant correlations of MEP and TEP amplitude per component and trial on group-level are depicted in **Figure A1** in the Appendix.

## Discussion

Within this study, the effect of the induced current direction on cortical and cortico-spinal excitation was investigated in 23 healthy subjects. By applying 200 biphasic pulses of opposite polarities at subject’s individual motor hotspot using a cobot-assisted TMS setup, MEPs, TEPs, LMFP and GMFP were contrasted between current directions (AP-PA vs. PA-AP). The results confirm a direction-specific excitation pattern in M1 for the RMT, MEP latency, P60 latency, N100 amplitude and GMFP as well as LMFP for mid-components (51-110 ms post TMS-pulse). However, no significant differences were found between current directions for MEP amplitude, other TEP component amplitudes and latencies, early- or late GMFP or LMFP. Significant positive correlations on group level were found for MEP amplitude and N100 in both current directions and for MEP amplitude and P30 component using AP-PA induced currents.

### Motor Evoked Potentials

As expected, we observed higher RMT and longer MEP latency with induced PA-AP currents in comparison to AP-PA but no difference in MEP amplitude. Our findings align with prior results regarding the directional sensitivity of M1 stimulation: the AP directed current flow in the second phase predominantly activated later I-waves, therefore requiring a higher stimulation intensity to reach threshold with increased duration of the descending pathway (Davila-Pérez et al., 2018; Delvendahl et al., 2014; Di Lazzaro et al., 2008; Di Lazzaro, Oliviero, Saturno, et al., 2001; Di Lazzaro & Rothwell, 2014; Sakai et al., 1997; Sommer et al., 2006; Weyh et al., 2005). In the specific case of biphasic stimulation, Davila-Pérez and colleagues argue that at suprathreshold level of stimulation, a contradictory effect of the antagonistic phases causes the prolonged latencies due to an imbalanced activation of excitatory and inhibitory inter-neuronal networks (Davila-Pérez et al., 2018), which is reflected by longer MEP latency observed for TMS with induced current in PA-AP direction in the present study. On a descriptive level, we also observed a lower MEP amplitude with AP-PA directed currents as previously described by Guidali and colleagues (2023). This could indicate that the cortico-spinal excitability, as measured by MEP amplitude, converges at the descending pathway when the stimulation is adjusted to threshold and found differences in the TEPs are not modulated by the threshold-adjusted stimulation intensity (Guidali et al., 2023). However, our results are descriptive and should be interpreted with caution. Since biphasic pulses are predominantly used in clinical treatment due to technical limitation of emitting more selective monophasic pulses in a repetitive protocol, further investigation of the cortical activation pattern is crucial.

### Transcranial Evoked Potentials

From all TEP components examined (N15, P30, N45, P60, N100, P180), significant differences between current directions were only found for the P60 latency and N100 amplitude. The average P60 peak occurred later with AP-PA orientated currents while the N100 peak was higher, i.e., more negative with PA-AP currents.

P60 and N100 are discussed to reflect reafferent feedback from the periphery. N100 could additionally reflect inhibitory neural activity or occur as a result of sensory processing or sensory contamination (Ahn & Fröhlich, 2021; Darmani & Ziemann, 2019; Schoisswohl et al., 2024). Since we did not find the same difference for the P180 component, which can also indicate late auditory processing (ter Braack et al., 2015) and should systematically vary in the same manner as the N100, the measured N100 amplitude is likely not modulated by auditory input but rather results from cortical activation via TMS. If we assume that N100 is an GABA-A and GABA-B receptor mediated inhibition marker, since its amplitude correlates with the length of the cortical silent period (Belardinelli et al., 2021; Darmani & Ziemann, 2019; Farzan et al., 2013), the PA-AP current direction could have induced higher cortical inhibition. Our results regarding the prolonged P60 latency when applying AP-PA currents is in contrast to the reduced latency of the MEP with this current direction. This is contrary to the assumption that P60 represents the reafferent feedback of the evoked motor response, since both the motor and transcranial evoked potentials should vary in the same manner.

Since we did not find differences between earlier components (N15-N45), initial cortical activation to stimulation seems stable across current directions with our experimental setup and pre-processing procedure and is in contrast to the results described by Guidali et al. (2023) who measured a smaller P15 component amplitude and shorter latency when using induced AP-PA currents in comparison to PA-AP currents or the results of Beck et al. (2024) who described a larger amplitude in the first peak and smaller amplitudes in the second and third peak of i-TEPs elicited with induced AP-PA currents. The observed differences in later components after 60 ms could reflect different processing pathways (Casula et al., 2018; Guidali et al., 2023).

Our results are partially in line but also contradictory to the results from Lucarelli et al. (2025). While describing strong influence of monophasic stimulation on TEPs over M1 due to the higher selectivity of the pulse current, alternating biphasic stimulation caused smaller TEP variability (Lucarelli et al., 2025). This is also described by Casula and colleagues (2018), who hypothesized that the reversed phase of a biphasic pulse at least partially could impede the activation of neurons involved in the origin of later potentials. In their comprehensive work, Lucarelli et al. (2025) describe significant directional-sensitivity of N15, P30, N45 and P180 amplitude as well as for N15 latency, elicited with the same stimulation intensity of 110% RMT as used in this study. However, they did not report any differences for P60 latency or N100 amplitude like we did (Lucarelli et al., 2025). The discrepancy in the results can only be explained speculatively: on the one hand, different pre-processing methods were used, for example 4 vs. 6 electrodes were defined for cluster identification per component or ICA was applied at different timepoints in the pre-processing pipeline. Other differences are, for example, the hardware used (e.g., MagVenture vs. Magstim stimulators with different coil diameters, 65 vs. 70 mm) or achieving the reversal of the current direction by rotating the coil to 180° vs. reversing the current flow through settings on the stimulator. It is possible that these differences led to different experimental outcomes, as described in the limitation section. However, as we found fewer differences in EEG responses, we can summarize by agreeing with the interpretation of Lucarelli and colleagues (2025) that the evoked transcranial potentials appear relatively stable across biphasic current directions.

Regarding global mean activation, our results are partially in line with previous studies: just as Casula and colleagues, who studied cortical responses in fronto-central regions, we did not observe any differences in metrics of global and local brain activity between applied current directions for early- and late responses. However, for the mid-responses, we observed the significantly higher amplitude corresponding to the timeframe of the N100 component in the M1 area, as discussed above. Since Casula et al. used a so-called controllable TMS device (cTMS) applying pulses of different shape and width, the global early- and late activation pattern can be considered as stable not only between used current directions but also across technical parameters pulse shape and width (Casula et al., 2018).

### Correlation of Motor- and Transcranial Evoked Potentials

Positive correlations between MEP amplitude and TEP amplitude were limited to the N100 component in both current directions and to the P30 component in the AP–PA current direction.

A similar correlation pattern on single-trial level, i.e. a trend towards higher MEP amplitude and associated N100 amplitudes was also observed by Roos and colleagues (2021). The authors concluded that the lack of significant correlation between MEP and TEP amplitude indicates TEP as not primarily or exclusively generated by M1 stimulation (Roos et al., 2021). Paus et al. (2001) described a significant correlation of N100 and absolute MEP amplitude. Fecchio and colleagues (2017) also used biphasic stimulation and report this pattern of positive EEG and EMG amplitude correlation and explained it by a higher afferent proprioceptive feedback, caused by the higher amplitude of the muscle activity (Fecchio et al., 2017). Alternatively, the higher N100 amplitude could be a results of muscle fibre depolarization after the muscle movement (Roos et al., 2021).

The P30-MEP correlation in the AP–PA condition could suggest direction- and intensity dependent activation at earlier steps of cortical pathways, i.e., without somatosensory feedback. Similar results on the N15-P30 complex and MEP amplitude were shown by Mäki & Ilmoniemi (2010) while applying monophasic pulses at threshold intensity. However, there are also contradictory results, e.g. Bonato et al. (2006) could not find any correlation between early components up to 30 ms post-pulse and MEP amplitude, applying pulses with the same suprathreshold intensity over M1 as we did. The authors argue that the absence of any correlations speaks in favour of the involvement of neuronal connections at the spinal level in particular in MEP genesis (Bonato et al., 2006). Despite a lack of clear behavioural correlates, TEPs are considered a more sensitive and direct marker of TMS-induced cortical excitability as TEPs also occur in the absence of MEPs (Casula et al., 2014; Roos et al., 2021; Thut & Pascual-Leone, 2010; Zhou et al., 2022).

In summary, significant correlations can be described both for early, direct activation components and for later components containing reafferent feedback with our experimental procedure. Given the partly contradictory results, further studies on correlations are needed.

### Limitations

For all results reported it is important to stress that these are specific for our experimental setup and preprocessing pipeline applied. It is known that the methodological choices influence the outcome of non-invasive brain stimulation studies (Beck et al., 2024; Brancaccio et al., 2024; Rogasch et al., 2022) and generalization of findings should take this into account. As example, we used a suprathreshold stimulation intensity (110% RMT) to elicit MEPs and TEPs at the same time. However, the different intensities per current direction can lead to differences in the EEG responses and limits the comparability with studies using subthreshold or even higher suprathreshold stimulation to focus on the cortical responses or saturation effects. The suprathreshold stimulation might have added somatosensory evoked potentials into the measured responses (Biabani et al., 2019), even while comprehensive noise masking was used. It is possible that a direct contact of the coil and the electrodes caused a bone conduction of sound and vibration since we did not cover it with an additional foam layer, leading to somatosensory co-stimulation and therefore a higher N100 amplitude. Also, the stimulation was not controlled in terms of the neural oscillations, preventing the stimulation from being linked to the underlying oscillatory phase of the motor cortex which influences early as well as late evoked potentials (Perera et al., 2024).

As with other TMS-EEG studies, due to the long duration of the experiment, it is likely that the subjects became tired and less attentive during the course of the study, despite the instructions not to fall asleep and to keep their eyes open. Noreika and colleagues (2020) demonstrated increase of MEP- and early TEP-amplitudes as well as variability with decreasing alertness levels. Finally, it is important to note that we have interpreted uncorrected *p*-values. After correcting for multiple comparisons via Bonferroni method, the only remaining significant contrasts are the current direction difference for the RMT and the correlation between MEP amplitude and N100 amplitude for both AP-PA and PA-AP currents.

## Conclusion

The present findings contribute to a growing body of TMS-EEG studies highlighting the importance of current direction in cortical and cortico-spinal responses to the stimulation and demonstrate the directional-sensitivity of the M1. Future studies should examine these direction-specific effects for their generalizability across larger samples, experimental setups and pharmacological interventions.

## Author Contributions

MO: Conceptualization, Data Curation, Methodology, Formal analysis, Investigation, Writing - Original Draft, Visualization

CK: Conceptualization, Data Curation, Methodology, Investigation, Writing - Review & Editing

WM: Resources, Writing - Review & Editing, Supervision, Project administration

BL: Resources, Writing - Review & Editing, Supervision

SSch: Conceptualization, Methodology, Writing - Review & Editing, Supervision, Project administration

## Acknowledgements

The study was funded by the dtec.bw – Digitalization and Technology Research Center of the Bundeswehr [MEXT project]. The dtec.bw is funded by the European Union – NextGenerationEU.

## Notes

### Competing Interest Statement

The authors have declared no competing interest.

