## Appendix for "Direction-Specific Effects of Biphasic Transcranial Magnetic Stimulation on Cortical and Cortico-spinal Excitability"

**Table A1**
*Results of EMG Measures and Comparison between Current Directions*

| | **AP-PA** | **PA-AP** | **Exact Wilcoxon signed-rank test** |
|---|---|---|---|
| RMT [%MSO] | 52.2 ± 5.6 | 60.3 ± 8.5 | $\boldsymbol{p \leq .001}$, $\boldsymbol{p_{adj} = .0017}$ |
| | *MD* = 52 | *MD* = 63 | CI [-10.500 -5.500] |
| Amplitude [µV] | 478.62 ± 386.24 | 578.05 ± 578.82 | $p = 0.7079$, $p_{adj} = 1$ |
| | *MD* = 350.03 | *MD* = 359.02 | CI [-127.699 60.395] |
| Latency [ms] | 24.64 ± 1.53 | 25.28 ± 1.43 | $\boldsymbol{p = 0.02901}$, $p_{adj} = .4352$ |
| | *MD* = 24.85 | *MD* = 25.2 | CI [-1.123 -0.0345] |

*Table A1*. Values depict the mean ± standard deviation or the median (MD). AP-PA: anterior-posterior to posterior-anterior current flow in the tissue. PA-AP: posterior-anterior to anterior-posterior current flow in the tissue. RMT: Resting motor threshold. CI: confidence interval. %MSO: percentage of maximal stimulator output. $p_{adj}$: $p$-value after multiple comparison adjustment.

**Table A2**

*Results of exact Wilcoxon signed-rank test comparisons of TEP peak amplitude and latencies between current directions*

| | **Amplitude [µV]** | **Latency [ms]** |
|---|---|---|
| | **AP-PA vs. PA-AP** | |
| N15 | $p$ = .5399 | $p$ = .5803 |
| | $p_{adj}$ = 1 | $p_{adj}$ = 1 |
| P30 | $p$ = .2467 | $p$ = .5803 |
| | $p_{adj}$ = 1 | $p_{adj}$ = 1 |
| N45 | $p$ = .41 | $p$ =.8932 |
| | $p_{adj}$ = 1 | $p_{adj}$ = 1 |
| P60 | $p$ = .8932 | ***p* =.03267*** |
| | $p_{adj}$ = 1 | $p_{adj}$ = .4900 |
| N100 | ***p* = .009146**** | $p$ =.8229 |
| | $p_{adj}$ = .1372 | $p_{adj}$ = 1 |
| P180 | $p$ = 0.3447 | $p$ = 0.6221 |
| | $p_{adj}$ = 1 | $p_{adj}$ = 1 |

*Table A2.* AP-PA: anterior-posterior to posterior-anterior current flow in the tissue. PA-AP: posterior-anterior to anterior-posterior current flow in the tissue. Significant p-values are highlighted with asterisks *. $p_{adj}$: $p$-value after multiple comparison adjustment.

### Correlation Analysis

#### Figure A1

*Correlation of MEP- and TEP amplitudes*

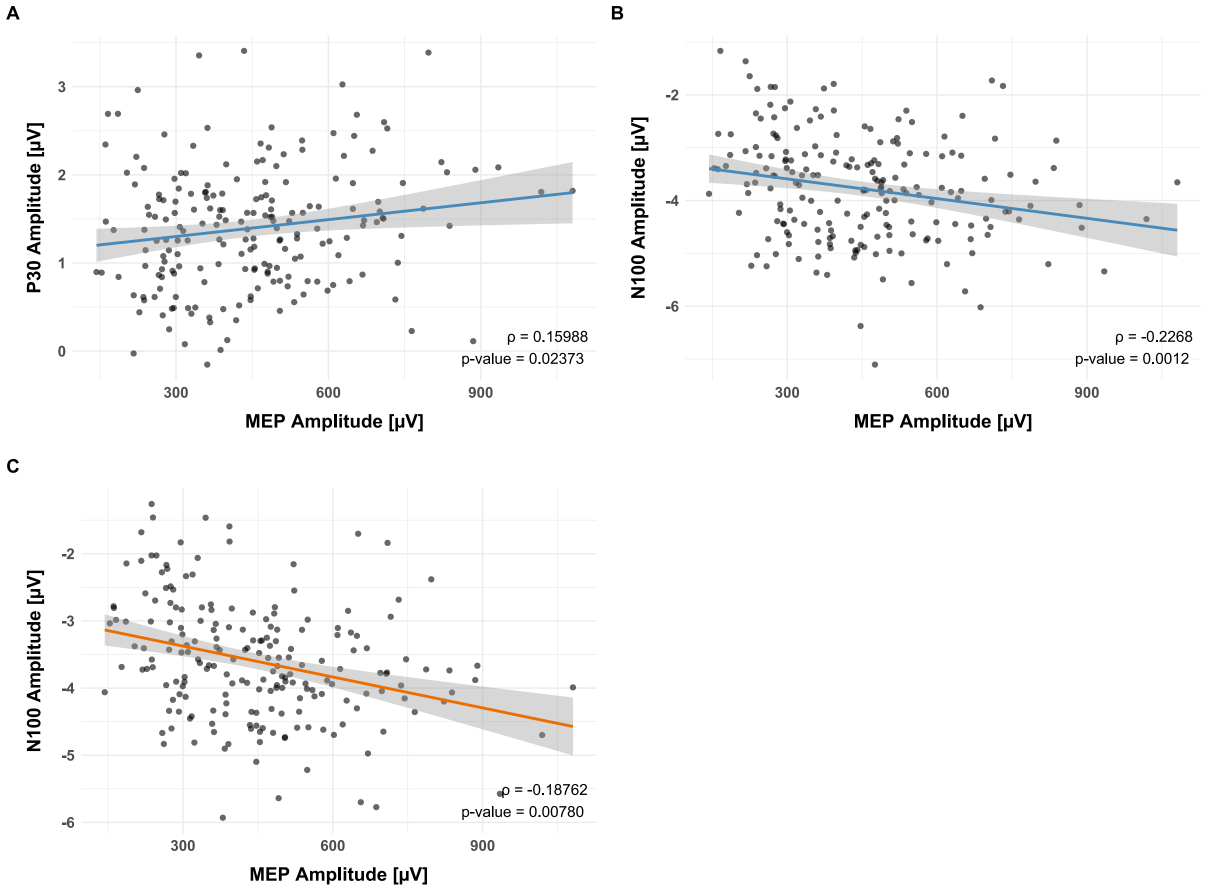


*Figure A1.* A: Significant Spearman correlation at group level between P30 TEP component amplitude and MEP amplitude. B: Significant Spearman correlation at group level between N100 TEP component amplitude and MEP amplitude. C: Significant Spearman correlation at group level between N100 TEP component amplitude and MEP amplitude. Blue line = AP-PA; orange line = PA-AP. ρ = Spearman's rank correlation coefficient, *p* = p-value. Bonferroni corrected *p*-values: P30 TEP component amplitude and MEP amplitude (A) $p_{adj}$ = .119; N100 TEP component amplitude and MEP amplitude (B) $p_{adj}$ = .0062; N100 TEP component amplitude and MEP amplitude (C) $p_{adj}$ = .0390.
